# A spontaneous mutation in a key C_4_ pathway gene significantly alters leaf δ^13^C, uncoupling its relationship with WUE and photosynthetic performance in *Zea mays*

**DOI:** 10.1101/2025.02.20.639358

**Authors:** Robert J. Twohey, Joseph D. Crawford, Matthew J. Runyon, Jiayang Xie, Andrew D. B. Leakey, Asaph B. Cousins, Anthony J. Studer

**Affiliations:** Department of Crop Sciences, University of Illinois Urbana-Champaign, Urbana, IL 61801, USA; School of Biological Sciences, Washington State University, Pullman, WA 99164, USA; Department of Plant Biology, University of Illinois at Urbana-Champaign, Urbana, Illinois 61801, USA; Institute for Genomic Biology, University of Illinois at Urbana-Champaign, Urbana, Illinois 61801, USA

**Keywords:** *Carbonic Anhydrase 1* (*cah1*), Leaf δ^13^C, QTL Mapping, Stomatal density, Water Use Efficiency, *Zea mays*

## Abstract

Increases in global temperature and drought are negatively impacting the yields of major crops. Therefore, targeted improvements to intrinsic water use efficiency (*WUE_i_*) are needed to reduce the water required for agricultural production. While it is very time-consuming to directly measure *WUE_i_,* stable carbon isotope ratios (δ^13^C) are a reliable high throughput proxy trait for quantifying *WUE_i_* in C_3_ species. While genetic studies have improved our understanding of the relationship between *WUE_i_* and δ^13^C in C_4_ species, the knowledge needed to implement δ^13^C in breeding schemes is incomplete. Using a *Zea mays* line with an extremely negative δ^13^C value, a quantitative genetics approach was used to identify a large deletion in *carbonic anhydrase1* (*cah1*). Carbonic anhydrase is the first enzymatic step of the C_4_ photosynthetic pathway and is known to affect δ^13^C. Surprisingly, the line with the mutant allele has significantly higher carbonic anhydrase activity with a concurrent reduction in δ^13^C, opposite of what would be expected based on C_4_ carbon isotope fractionation theory. These observed decouple δ^13^C and *WUE*_i_, which calls for further investigation into carbon isotope discrimination in C_4_ species.

## Introduction

Changes in global temperature and precipitation patterns present significant challenges to the sustainable crop production required to support the feed, fuel, and fiber needs of the growing human population. To combat environmental stress, new technologies and practices are needed that increase the climate resiliency and productivity of major crop species (Eckardt *et al.,* 2023). The global surface temperature has risen to 1.1°C above preindustrial levels and models predict that trend to continue, reaching 1.5°C by 2040 (IPCC, 2022). In addition to increasing atmospheric temperatures, precipitation patterns have become more sporadic causing droughts to become more intense and frequent (Gamelin *et al.,* 2022). Increased drought sensitivity and reductions in crop yield have been observed as a result (Lobell *et al.,* 2014; Cooper and Messina, 2023). Over the past 20 years, drought-induced agricultural losses have surpassed one billion dollars (Leeper *et al*. 2022). These events highlight the need to develop plants that can delay the onset of drought stress by using less water while achieving equivalent or greater production i.e. increasing water use efficiency (*WUE*) (Leakey et al. 2019).

Under drought conditions, plants become water deficient and require supplemental water to remain productive. Increasing irrigation to overcome water limitations is a dangerous strategy given that 70% of global freshwater use is already attributed to agriculture (McDaniel *et al.,* 2017). Therefore, increasing crop productivity under challenging growing conditions without increasing water use is necessary to avoid depleting freshwater reserves. One target trait that has the potential to improve crop resilience and sustainability is *WUE*, defined as the amount of carbon assimilation relative to water used by the plant (Vadez *et al.,* 2014; Ellsworth and Cousins, 2016; Leakey et al., 2019). Agronomic *WUE* (*WUE_a_* = yield/transpired water) defines *WUE* at a whole plant level in terms of production, while intrinsic *WUE* (*WUE*_i_ = net photosynthesis/stomatal conductance) measures instantaneous efficiency at the leaf level (Dhanapal *et al.,* 2015; Ellsworth and Cousins, 2016). Selection for *WUE*_i_ improvements cannot be at the expense of net CO_2_ assimilation, otherwise reductions in productivity will likely be observed (Leakey *et al.,* 2019).

The identification and development of a commercial variety with improved *WUE*_i_ can be very time-consuming. Current methods of phenotyping *WUE*_i_, such as gas exchange experiments, limit the number of genotypes that can be screened. Therefore, there is a need for an alternative high-throughput method to measure *WUE*_i_. Leaf tissue stable carbon isotope composition (δ^13^C_leaf_) is an integrated measurement of CO_2_ availability and carbon metabolism potentially providing a proxy for the relationship between stomatal conductance and net CO_2_ assimilation that can be scaled up to breeding programs easier than measurements of leaf gas exchange (O’Leary, 1981; Condon *et al.,* 2004). Since stomatal conductance regulates CO_2_ uptake and transpiration, theory states that changes in *WUE*_i_ should ultimately affect δ^13^C_leaf_ (Farquhar and Richards, 1984). The use of δ^13^C_leaf_ for screening *WUE*_i_ in large breeding programs has shown promise in C_3_ species (Farquhar and Richards, 1984; Rebetzke *et al.,* 2002; Teulat *et al.,* 2002; Condon *et al.,* 2004; Saranga *et al.,* 2004; Chen *et al.,* 2011). While the relationship between *WUE*_i_ and δ^13^C_leaf_ is often observed and straightforward to interpret in C_3_ species, the carbon-concentrating mechanism in the C_4_ photosynthetic pathway makes the relationship between δ^13^C_leaf_ and *WUE*_i_ more complex (Cernusak *et al.,* 2013). The ratio of intercellular to ambient CO_2_ partial pressure (*C*_i_/*C*_a_), phosphoenol pyruvate carboxylase (PEPC) discrimination, and Rubisco discrimination all effect δ^13^C_leaf_ (Farquhar, 1983; von Caemmerer *et al.,* 2014). In addition to the major enzymes that fractionate carbon isotopes, δ^13^C_leaf_ in C_4_ species is also influenced by the loss of CO_2_ that was fixed in the form of C_4_ acids and decarboxylated in the location of Rubisco but ultimately leaking out before it could be fixed by Rubisco, referred to as leakiness (Φ) (Kromdijk *et al.,* 2014). However, recent studies have revealed that the overall contribution of Φ to differences in δ^13^C_leaf_ remains consistent in C_4_ species in response to growth under changing environmental conditions (Henderson *et al.,* 1992; Sonawane and Cousins, 2020). The influence of post-photosynthetic fractionation on δ^13^C_leaf_ also needs to be considered but differences in its contribution is likely small within a species (Tcherkez *et al.,* 2011).

However, further studies are still necessary to identify and improve the understanding of underlying mechanisms controlling photosynthetic and post-photosynthetic discrimination in diverse C_4_ plants before δ^13^C_leaf_ can be implemented as a proxy of *WUE_i_*.

Identifying genetic regulators controlling δ^13^C_leaf_ has significant potential to improve *WUE* and reduce the environmental footprint of agriculture. *Zea mays* holds great economic importance as a major crop (an estimated 94.1 million acres planted in the USA in 2022, USDA-NASS). The large amount of genetic diversity and availability of genetic resources also makes *Z. mays* an ideal model organism to study water use in a C_4_ crop (Buckler and Stevens, 2006). Recent studies have standardized methods of leaf isotope sampling and achieved a better understanding of the relationship between δ^13^C_leaf_ and water use traits in *Z. mays* (Kolbe et al., 2018; Avramova *et al.,* 2019; Twohey III *et al.,* 2019; Sorgini *et al.,* 2020; Eggels *et al.,* 2021; Crawford *et al.,* 2023). However, only a small number of studies have identified genes that alter δ^13^C. A candidate gene, ZmAbh4, controlling kernel isotopic composition was identified using genetic mapping approaches (Gresset *et al.,* 2014; Avramova *et al.,* 2019). ZmAbh4 was later targeted using CRISPR and mutated allelic variants were shown to alter kernel ^13^C isotope discrimination (Δ^13^C) and plant *WUE* traits (Blankenagel *et al.,* 2022). Future discoveries could further elucidate other genes in the ABA pathway, or novel pathways involved in water use and CO_2_ assimilation. While these studies support the efficacy of using δ^13^C as a high throughput proxy for *WUE*_i_, further understanding of δ^13^C genetic control in *Z. mays* is required before it can be implemented into breeding strategies.

Our study investigates an elite *Z. mays* line showing a relatively negative δ^13^C_leaf_ signature to elucidate genetic mechanisms altering δ^13^C_leaf_ and improve our understanding of the relationship between δ^13^C_leaf_ and *WUE* in a major C_4_ crop. Initial biparental mapping identified a large QTL correlated with the observed δ^13^C_leaf_ signature. The significant QTL interval contained a large list of candidate genes; therefore, a targeted genetic complementation test and sequencing approach was used to narrow down the region of interest and identify the mutation driving the observed δ^13^C_leaf_ signature. Further investigation of this unique line included physiological and molecular characterization to identify the causative mutation and tease apart the relationship between photosynthetic performance, *WUE*, and δ^13^C_leaf_.

## Materials and Methods

### Plant Growth

The ExPVP lines OQ414 (PI 583776) and LH82 (PI 601170) were ordered from the USDA Germplasm Recourses Information Network (GRIN) and are publicly available. The two lines were selfed during their first planting to make stocks of seed for future experiments.

During the 2018 and 2019 field seasons, a panel of 393 ExPVP lines, including OQ414 and LH82, was planted in two field locations at the Crop Sciences Research and Education Center located in Urbana, Illinois, USA. The two locations are approximately 1.6km apart. One location was irrigated while the other was rain-fed. Twenty kernels were planted in a 3.7m row with 0.8m spacing between rows and 0.9m alleys. OQ414 and LH82 were replicated multiple times in a randomized block design.

During the winter of 2019, OQ414 and LH82 were crossed in the first greenhouse generation and F_1_ individuals were again selfed during the second greenhouse generation to produce the F_2_ mapping individuals. In 2020, the F_2_ population was grown in Champaign County at a private farm 32.2 km away from the University of Illinois. Row spacing was the same as described above.

The genetic complementation test was grown at the University of Illinois Plant Care Facility greenhouses located in Urbana, Illinois, USA. Kernels were planted in a 50-cell tray and 5 individuals of each genotype were transplanted into Classic 600 pots two weeks after germination. A 3:1:1:1 soil mix of BM6, sterilized field soil, peat, and perlite was used. Growing conditions included LED supplemental light for 14-h days and day/night temperatures were set at 28/24°C. After the transplant, each individual pot was fertilized once with Sprint 330 Chelated Iron. The plants also received approximately 1 liter of 3 ppm CalMag (15-5-15) every 7 days.

Tissue samples were collected from the uppermost fully expanded leaf once the plants reached V10. Leaf tissue collection is described below.

### δ^13^C_leaf_ Tissue Collection

All δ^13^C_leaf_ sample collection was performed as described in Twohey III *et al*. (2019). During the 2018, 2019, and 2020 field seasons, δ^13^C_leaf_ samples were collected from healthy, uppermost fully expanded, V10 leaves. Depending on the experiment, leaf samples were collected as pooled samples from a row or individual plants. The 2018 and 2019 isotope collections were collected as pooled samples. Six leaf punches were collected from four separate plants in the same row of the field. All other experiments, where isotopes were collected, consisted of isotope samples from individual plants. Each sample contained 6-12 tissue punches from each side of the midrib totaling 12-24 leaf punches per sample. Leaf tissue samples were then dried at 65°C for 7 days, ground, and stored in a desiccation cabinet until isotope analysis.

### Stable Carbon Isotope Analysis

Leaf stable carbon isotope analysis was run as described in Twohey III *et al*. (2019). The samples were run through a Costech Instruments elemental combustion system, and then a mass spectrometer to determine δ^13^C values. The spectrometer used varied based on run location.

Samples processed at Washington State University were run on a Delta PlusXP and University of Illinois samples were on a Delta V Advantage. The precision of both instruments has been shown to be ±0.2‰ when measuring δ^13^C. The 2018 and 2019 δ^13^C_leaf_ field samples and kernel δ^13^C samples were processed at the University of Illinois. All other samples were processed at Washington State University. Vienna Peedee Belemnite was used as a calibration standard at each facility, insuring consistency across all data.

### GBS Library

Leaf tissue was collected from 341 F_2_ individuals and four replicates of each parent (OQ414 and LH82) grown during the 2020 field season. High-quality DNA was extracted from the tissue samples using a modified CTAB extraction protocol with double chloroform:isoamyl alcohol 24:1 and 70% ethanol washes. DNA concentrations were normalized using PicoGreen quantification (Invitrogen, Carlsbad, CA). A revised genotype-by-sequencing library protocol was used from Elshire *et al*. (2011). Genomic DNA was digested using a double restriction digest containing PstI and ApeKI enzymes. GBS libraries were constructed in 96 well plates (VWR 89049-178). In each well, 100ng of DNA, 1.5uL of 0.1mM rare PstI barcoded adapter, and 0.5 uL of 10uM common barcoded adapter were added. One pooled library, containing all samples, was run through a size selection protocol using Agencourt AMPure XP Beads (Beckman Coulter Inc., Brea, CA). The size selected library was amplified for 15 cycles using KAPA HiFi HotStart ReadyMix (KAPA Biosystems), and then a final size purification using Agencourt AMPure XP Beads was completed. Using PicoGreen, a 1ng/uL dilution of the library was made using 10 mM Tris and 5uL of the dilution was run on a 1% agarose gel to verify fragment size. A Bioanalyzer 2100 (Agilent, Santa Clara, CA) was used to estimate fragment sizes in the pooled library before sequencing. The library was diluted to 10nmol using 10 mM Tris and was run on two NovaSeq6000 SP lanes using 100-bp single-end reads (Roy J. Carver Biotechnology Center, University of Illinois, Champaign, IL, USA). The Tassel 5 GBS v2 Pipeline and B73v5 reference genome were used to identify SNPs (Glaublitz *et al.,* 2014).

Before using the SNP data for mapping, multiple filtering options were applied. All individuals containing 25% or more missing data across all SNPs were removed. Any markers that had an average coverage of 40% or less across all individuals were removed. Heterozygosity (0.4 to 0.6) and minor allele frequency (0.46 to 0.6) ranges were used to further filter the marker set. The final SNP set consisted of 1,414 markers and 274 individuals including the parents.

### QTL Mapping

The QTL mapping was executed using the R package (rqtl) (Broman *et al.,* 2003). The functions “calc.genoprob” and “est.map” were used to make a genetic map. The Kosambi function was run with a 0.01 error probability. Using “scanone”, QTL were identified across the genome. A total of 10,000 permutations were used to determine the LOD threshold cutoffs. The “makeqtl”, “fitqtl”, and “refineqtl” functions were used to identify significant QTL while using the Haley-Knott Regression method. This process was repeated until no new significant QTL could be added to the model using a 0.01 significance threshold. The final 1 LOD mapping intervals were calculated using “lodint”.

### Stomatal Imaging

Additional leaf tissue samples were collected from the F_2_ population grown in 2020 to measure stomatal traits. A leaf tissue sample of approximately 1.5 x 0.5 cm was collected from the center of the same leaf as isotope samples. The sample was placed in a 2ml tube and immediately frozen in liquid nitrogen. The tissue samples were attached to a microscope slide and imaged using a Nanofocus μsurf Explorer Optical Topometer (Oberhausen, Germany) at 20X magnification with 0.6 numerical aperture as described in Xie *et al*. (2021). Each tissue sample was imaged 3 times on each side of the leaf (adaxial and abaxial) for a total of 6 images collected from each F_2_ plant.

Stomata were automatically counted using a Mask R-CNN model (Xie *et al*., 2021). 100 images were randomly selected for ground-truthing, as recommended by Tan *et al*. (2024). These manual counts indicated that the accuracy and recall of the ML was almost perfect, at 0.972 and 0.999, respectively. The entire data set of stomatal images was then run through the model and total stomatal counts were used to QTL map stomatal density in the F2 mapping individuals as described above in *QTL Mapping*.

### OQ414 gDNA and cDNA Sequencing

The first approach to identify a causative mutation in OQ414 was through Sanger sequencing completed by the Roy J. Carver Biotechnology Center located at the University of Illinois Urbana-Champaign. Using OQ414 and LH82 gDNA, polymerase chain reaction (PCR) was used to amplify portions of the *cah1* gene (also previously referred to as *ca1*). Using 3:220993296/3:220987483 and 3:220992441/3:220988556 (Supp.1) primer pairs, *cah1* fragments were amplified and sequenced from the forward and reverse primers. After the *cah1* mutation as located in OQ414, Whole gene linear sequencing was performed by Plasmidsaurus using Oxford Nanopore Technology with custom analysis and annotation paired with additonal Sanger sequencing by the Roy J. Carver Biotechnology Center. Using 3:220994897/3:220993218, 3:220992158/3:220993296, 3:220992441/3:220988556, 3:220988556/3:220987483, 3:220992441/3:220988451, and 3:220989065/3:220988451 primer pairs, *cah1* fragments of LH82 and OQ414 were amplified through PCR, purified, and sequenced.

A trizol RNA extraction protocol and SuperScript VILO reverse transcriptase kit was used to make OQ414 and LH82 cDNA. Then OQ414 cDNA was amplified through PCR using primers 3:220994895/3:220987666 (Supp.1) to target the *cah1* gene from exon 1 through the 3’UTR. The OQ414 PCR product was purified, and a Zero Blunt TOPO PCR Cloning Kit was used to isolate splice variants of OQ414 cDNA. The five largest OQ414 cDNA plasmids were submitted for sequencing and then aligned to the B73v5 reference sequence. LH82 cDNA was sequenced using 3:220994897/3:220989560, 3:220990512/3:220987793, 3:220992974/3:220992393, and 3:220994897/3:220992933. Whole plasmid sequencing was performed by Plasmidsaurus using Oxford Nanopore Technology with custom analysis and annotation. Linear Sanger sequencing was performed by the Roy J. Carver Biotechnology Center. Protein amino acid sequence was estimated for LH82 and the cDNA OQ414 splice variants using the Expasy- Translate tool (https://web.expasy.org/translate/) and then aligned in MEGA11: Molecular Evolutionary Genetics Analysis version 11 (Tamura, Stecher, and Kumar 2021). The Muscle algorithm was used for protein alignment (Edgar, 2004).

### Gas exchange Measurements

Growth chamber grown plants were measured on the uppermost fully expanded leaf using a LI- 6800 (LI-COR Biosciences, Lincoln, NE) with a 3 cm^2^ square light source. The flow rate was 250 µmol m^-2^ s^-1^. The light intensity was 2,000 µmol m^-2^ s^-1^ with 90% red and 10% blue light quality. The relative humidity of the leaf chamber was set to 70%. The temperature of the leaf was set to 30 degrees Celsius. The initial partial pressure of CO_2_ was 33 Pa (400 ppm). Leaf measurements were recorded every 5 sec. After the leaf acclimated for ∼ 20 minutes to conditions, an autoprogram was started to record observations at the same conditions for 4 min. The steady state gas exchange data presented is extracted from the first 4 min before conditions changed and averaged for N = 6 biological replicates.

To measure *A/C_i_* curves, CO2 concentration started at 400 μmol mol^−1^ and then changed sequentially: 0, 10, 25, 35, 50, 75, 115, 150, 200, 250, 300, 350, 400, 600, 750, 400 μmol mol^−1^.

The maximum carboxylation capacity of phosphoenolpyruvate (*V*_pmax_) was calculated using the initial slope of the *A*/*c*_i_ curve (von Caemmerer, 2000), and CO_2_ saturated photosynthetic capacity (*A*_sat_) was estimated according to the horizontal asymptote of the *A*/*C*_i_ curve by using a four-parameter non-rectangular hyperbolic function as described in Markelz *et al*. (2011).

### PEPC, CA, and Rubisco Assays

Leaf discs were taken from the same growth chamber grown plants used for gas exchange measurements. The samples were immediately frozen using liquid nitrogen and then stored at - 80°C. The tissue was ground and aliquots were made as described in Crawford and Cousins, (2022). Quantification and analysis of CA, PEPC, and Rubisco is described in Studer *et al*. (2014).

## Results

### Identification of an extreme δ^13^C_leaf_ outlier in field-grown Zea mays

An Expired Plant Variety Protection Act (ExPVP) panel containing 393 inbred *Z. mays* lines was grown at the University of Illinois during the 2018 and 2019 field seasons. During each grow-out, one line (OQ414) was consistently identified to have an extreme δ^13^C_leaf_. During these trials, OQ414 had an average δ^13^C_leaf_ value of -14.35 ‰ ± 0.19 (2018) and -14.38 ‰ ± 0.2 (2019) (Table 1). Another ExPVP line (LH82) was selected from the panel for comparison because it has similar morphological characteristics to OQ414, but a distinctly different isotopic signature. LH82 had an average δ^13^C_leaf_ value of -11.76 ‰ ± 0.21 (2018) and -12.02 ‰ ± 0.17 (2019) (Table 1). These lines were grown again during the 2020 and 2021 field seasons, in which OQ414 consistently showed an extreme δ^13^C_leaf_ value of -14.84 ‰ ± 0.16 and -14.61 ‰ ± 0.13 respectively. The comparison line LH82 had δ^13^C_leaf_ values of -12.27 ‰ ± 0.19 and -12.41 ‰ ± 0.13 during 2020 and 2021 (Table 1). Every year these two lines were grown, OQ414 showed a significantly more negative δ^13^C_leaf_ value compared to LH82 (*T-test, p* < 0.01). Kernel δ^13^C was also measured from OQ414 and LH82 in 2020 to determine if the observed δ^13^C_leaf_ signature was reflected in the mature kernels. OQ414 and LH82 kernel δ^13^C values were similar to the observed δ^13^C_leaf_ values sampled from the same environment. Kernel δ^13^C was significantly different between OQ414 and LH82 (*T-test, p* < 0.0001), and averaged -14.29 ‰ ± 0.13 and -

**Table 1.**

| Year | LH82 | OQ414 | *p* value |
| --- | --- | --- | --- |
| 2018 | -11.76±0.21 | -14.35±0.19 | < 0.001 |
| 2019 | -12.02±0.17 | -14.38±0.20 | 0.01 |
| 2020 | -12.27±0.19 | -14.84±0.16 | < 0.0001 |
| 2021 | -12.41±0.13 | -14.61±0.13 | < 0.01 |

Presented as average ± standard error. Statistical significance was calculated within each year (*T-test*).

11.27 ‰ ± 0.03 respectively. Given these dramatic and consistent differences in δ^13^C_leaf_, OQ414 and LH82 were used to make a bi-parental mapping population to identify the genetic mechanism(s) controlling the extreme δ^13^C_leaf_ observed in OQ414.

### δ^13^C_leaf_ QTL Mapping

A bi-parental F_2_ mapping population was generated using OQ414 and LH82. The parent lines grown as controls with the F_2_ population had significantly different δ^13^C_leaf_ values (*T-test*, *p* < 0.0001). The δ^13^C_leaf_ analysis of the 314 F_2_ individuals showed a segregation ratio of 254:60 (4.3:1); which is close to a 3:1 ratio that would suggest δ^13^C_leaf_ differences in OQ414 and LH82 are controlled by a single gene with simple dominance/recessive alleles (Fig.1). Using genotype by sequencing (GBS) data (Supp.2), a genetic map was constructed. The genetic map contained an average of 141 ± 22 markers on each chromosome, with an overall average spacing of 3.8 cM between each marker. Haley-Knott regression identified two significant δ^13^C_leaf_ QTL peaks on Chr. 3 (Supp. 3). The first QTL contained a 1 LOD interval ranging from 464.1 cM to 489.1 cM (216.7 Mb to 223.2 Mb) with a peak marker located at 473.2 cM (218.1 Mb). The second QTL was identified approximately 6.7 Mb away containing a 1 LOD interval ranging from 290.6 cM to 302.9 cM (223.5 Mb to 225 Mb) with a peak marker located at 295.25 cM (224.7 Mb). To increase mapping resolution, additional markers were designed and tested within the region on Chr. 3 containing the two QTL intervals. An indel marker at position 3:222755101 (Supp. 1) showed a clear difference between the parental backgrounds and was used to genotype the F_2_ population. The marker was added to the QTL analysis and Haley-Knott regression identified a single significant δ^13^C_leaf_ QTL on Chr. 3 (Fig. 2). The QTL contained a 1 LOD interval ranging from 472.9 cM to 489.9 cM (218.1 Mb to 223.2 Mb) with a peak marker located at 482.4 cM (222.8 Mb) (Supp. 4). This position was consistent with the large effect QTL identified in the initial mapping.

**Figure 1:**
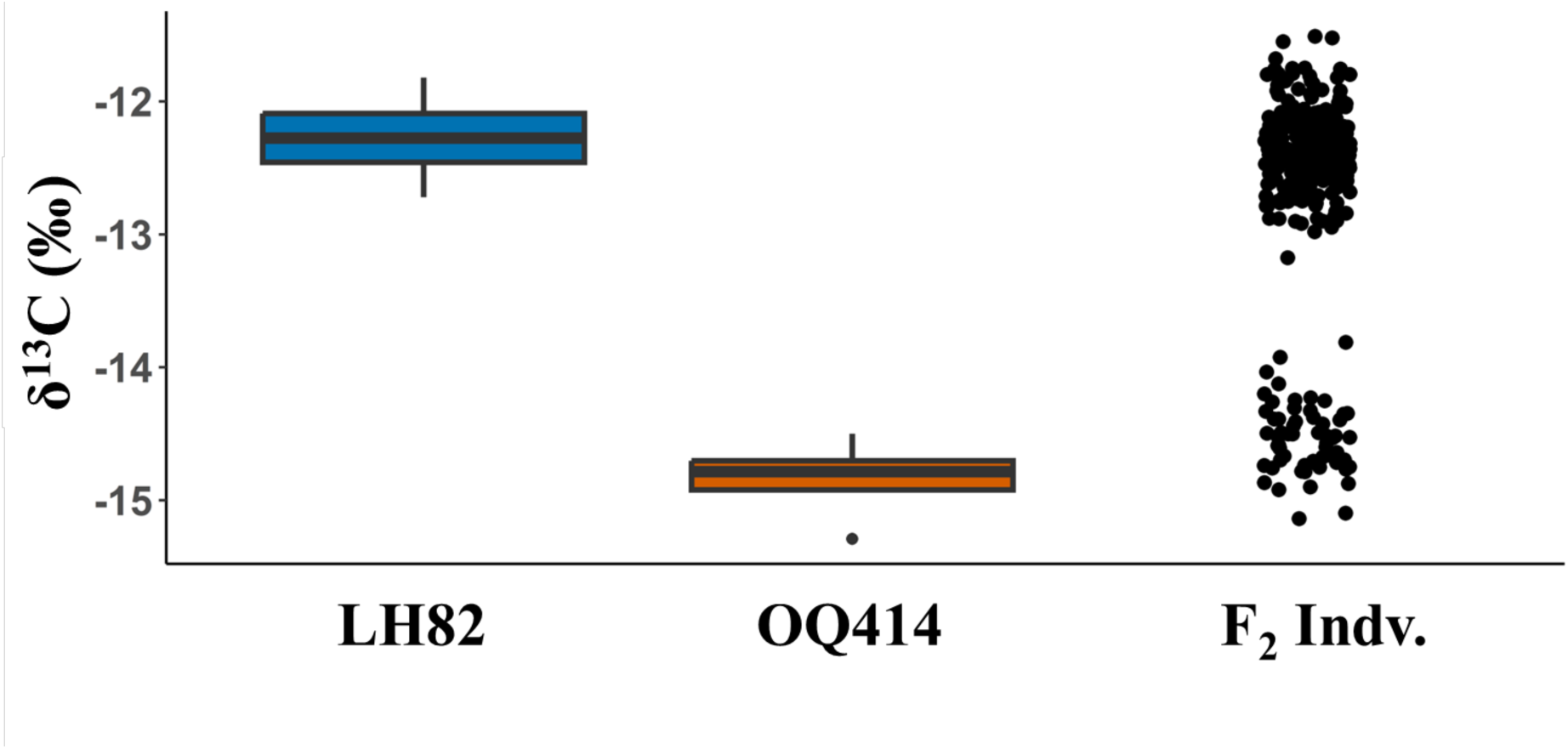
δ^13^C_leaf_ values of the F_2_ mapping population (black) and the two parental lines OQ414 (orange) and LH82 (blue) (n=4). OQ414 and LH82 have significantly different δ^13^C_leaf_ values (*T-test, p* < 0.0001). The black F_2_ Indv. circles represent 314 unique individuals. The box plots show the first and third quartiles, with middle horizontal lines indicating the median and the whiskers showing the mild outliers. Boxplot circles represent extreme outliers.

**Figure 2:**
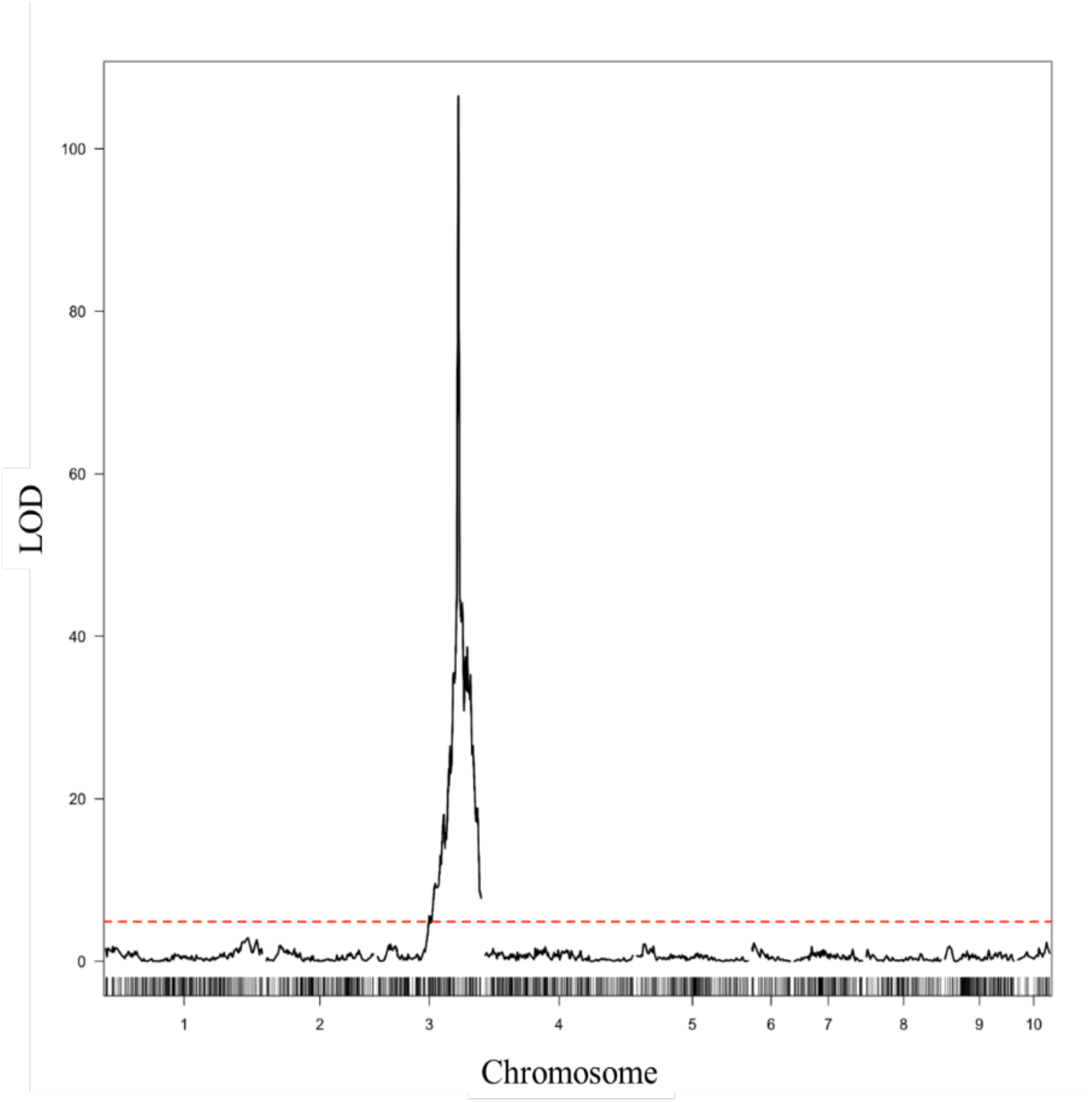
δ^13^C_leaf_ Haley-Knott Regression QTL Mapping. One δ^13^C_leaf_ QTL was identified in the F_2_ mapping population originating from the parents OQ414 and LH82. All 10 *Z. mays* chromosomes are represented on the X-axis. A significance threshold (red dashed horizontal line) was determined by 10,000 permutations and an alpha of 0.01.

### Stomatal Density QTL Mapping

Given the connection between stomatal conductance and δ^13^C, stomatal density data was also collected from individuals from the F_2_ mapping population. Haley-Knott regression identified multiple significant QTL for three stomatal density traits (abaxial density, adaxial density, and total stomatal density) on Chr. 4 and 5 (Fig.3, Supp. 4). Interestingly, Chr. 5 contained a significant QTL (peak, 394.9 cM) with overlapping 1 LOD intervals for abaxial and total stomatal densities. Adaxial density also showed an overlapping Chr. 5 peak but was not significant at a 0.01 cutoff (*p* < 0.013). No QTL colocalized for δ^13^C_leaf_ and the stomatal density traits, indicating that stomatal density was not the major mechanism driving the extreme δ^13^C_leaf_ values observed in OQ414.

**Figure 3:**
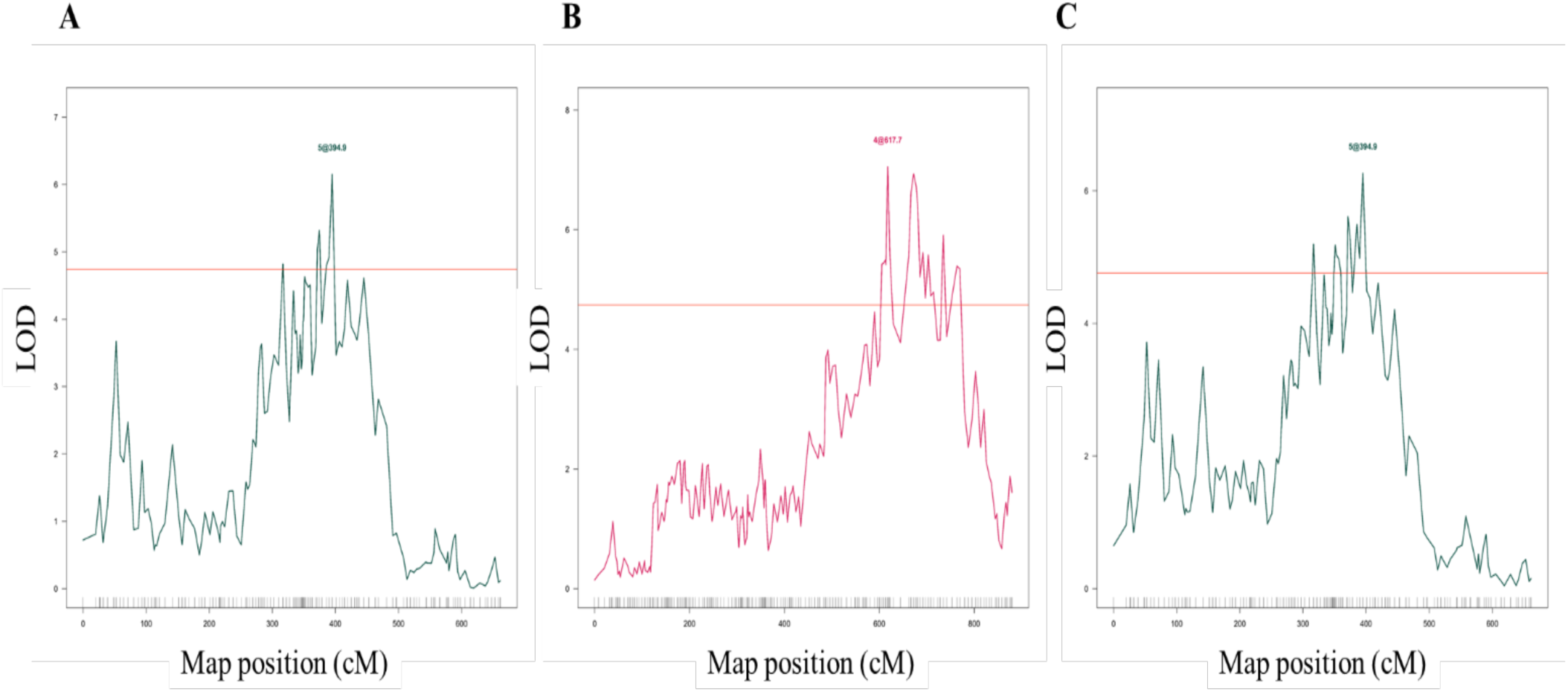
Stomatal density Haley-Knott Regression QTL Mapping. Stomatal density QTL were identified in the F_2_ mapping population originating from the parents OQ414 and LH82. Labels above each significant peak represent chromosome number and cM position. A) Abaxial stomatal density, B) Adaxial stomatal density, C) Total stomatal density (Abaxial + Adaxial). Significance threshold (solid red horizontal line) was determined by 10,000 permutations and an alpha of 0.01.

### cah1 Complementation Test

Previously reported δ^13^C_leaf_ values from maize carbonic anhydrase mutants showed negative δ^13^C values similar to OQ414 (Studer *et al*. 2014). Since carbonic anhydrase is a significant source of stable isotope fractionation in the C_4_ pathway, and given that the *cah1* and *cah2* genes are located within the 1 LOD interval of the major QTL on Chr. 3, we used a genetic complementation test to determine if the OQ414 δ^13^C_leaf_ phenotype is due to differences in either of the *ca* genes.

During the 2021 field season, OQ414 was crossed to *cah1*, *cah2,* and *cah1cah2* mutants. All three of the resulting F_1_ combinations were grown in the greenhouse with OQ414 and the *cah1cah2* mutant. By bringing the mutant *ca* alleles into the same genome as the OQ414 causative allele through crossing, the presence of the extreme δ^13^C_leaf_ phenotype in the resulting F_1_ would suggest that the mutation altering δ^13^C_leaf_ in OQ414 is in the same gene as the mutant *ca* allele. All F_1_ combinations containing the *cah1* mutant allele failed to complement and showed δ^13^C_leaf_ values lower than -13.3 ‰, while hybrids between OQ414 and *cah2* had normal δ^13^C_leaf_ values (Fig.4). This result suggests that OQ414 contains a functional *cah2* allele that rescues the *cah2* mutant phenotype. Since OQ414 could not complement the *cah1* mutant allele, a mutation in the OQ414 allele of *cah1* is likely causing the observed δ^13^C_leaf_ phenotype (Fig.4).

**Figure 4:**
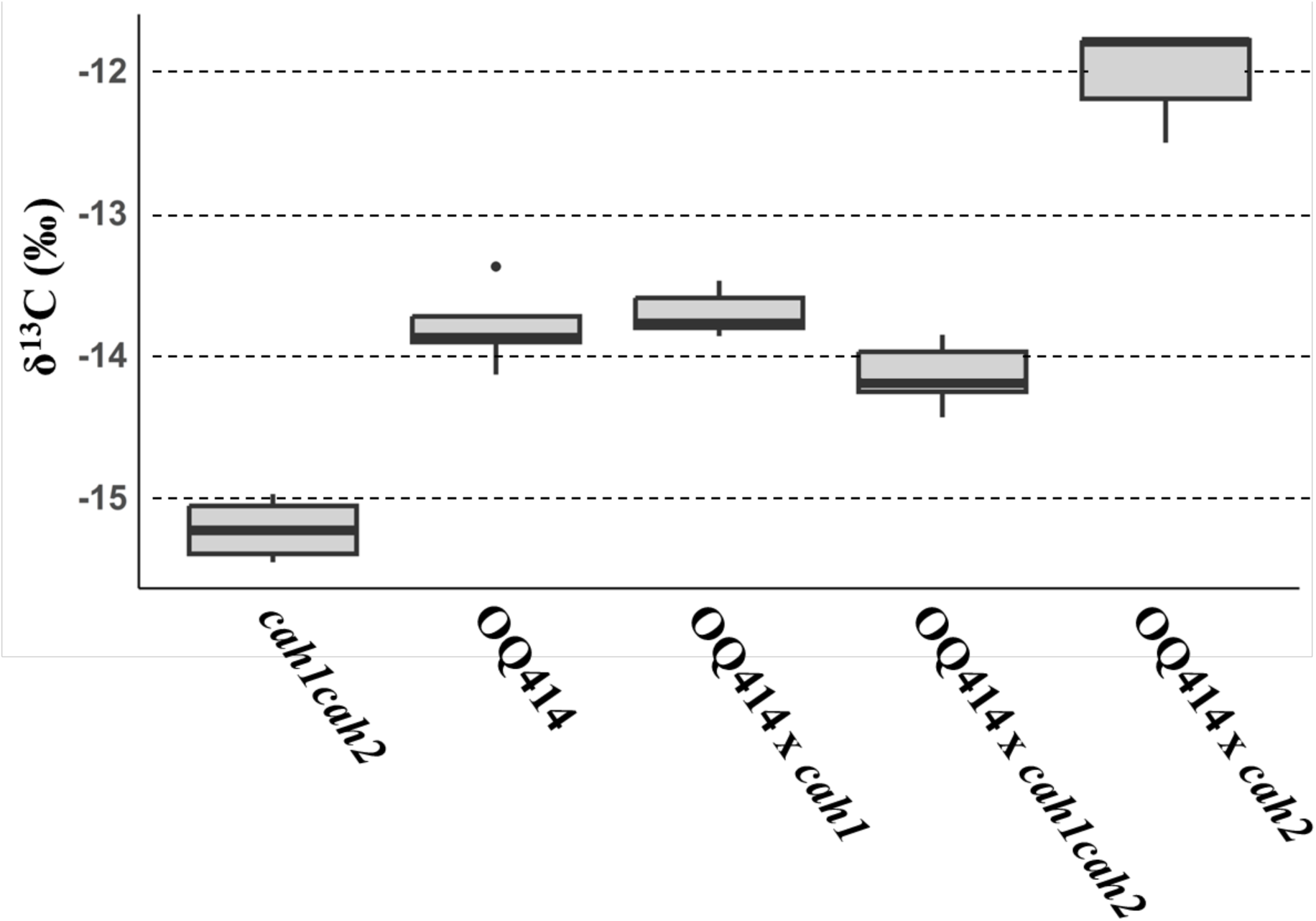
δ^13^C_leaf_ values from the greenhouse-grown complementation test. *cah1cah2* δ^13^C_leaf_ is significantly more negative than all other genotypes and OQ414 x *cah2* δ^13^C_leaf_ is significantly less negative than all other genotypes (ANOVA, p< 0.0001). The box plots show the first and third quartiles, with middle horizontal lines indicating the median and the whiskers showing the mild outliers. Box plot circles represent extreme outliers.

### OQ414 Causative Mutation in Carbonic Anhydrase

After the complementation test identified *cah1* as a candidate gene, a sequencing approach was used to identify the causal mutation in OQ414. Primers were used to PCR amplify the *cah1* gene and amplicons were Sanger sequenced. From this data a genomic DNA sequence of the OQ414 and LH82 alleles of *cah1* could be assembled (Supp. 5). After OQ414 and LH82 were aligned to the B73v5 reference sequence, a 3,191 bp deletion was identified in OQ414. The deletion removed exons 5-10 and the first 54 bp of exon 11 (Fig.5). To investigate the functional consequence of this deletion on the *cah1* transcript, OQ414 cDNA was produced in an RT-PCR reaction and multiple splice variants were identified (Supp.6). The sequenced transcripts confirmed the location of the OQ414 deletion in *cah1*, with all isoforms encoding the first 89 amino acids preceding the deletion (Supp.7). A transcript was recovered that retained the wild type coding sequence flanking the deletion except for 6 bp directly after the deletion in exon 11 (OQ10 and OQ16). Other splice isoforms were more variable and contained fragments of introns that resulted in frame shifts. cDNA splice variants OQ8 and OQ7 maintained 169 bp while OQ13 retains 71 bp of intron 4 sequence prior to the deletion. Variant OQ8 also retained intron 11 in the mature transcript. It is not known if these isoforms all produce proteins, but several contain long 3’UTRs that may result in nonsense mediated decay. Thus, the transcript resulting in a CA protein 222 amino acid shorter than wild type is likely the functional protein with altered fractionation of CO_2_.

**Figure 5:**
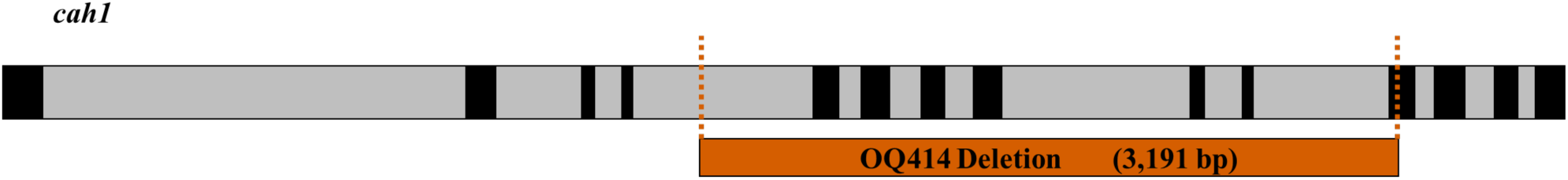
*cah1* (Zm00001eb158810) contains 14 exons and 13 introns. The orange rectangle and vertical dashed lines indicate the deletion in OQ414. This deletion removes the DNA sequence between a partial intron 4 and partial exon 11.

### Gas Exchange Physiology and Biochemistry of OQ414

Gas exchange measurements collected to determine steady state performance in the OQ414 and LH82 lines. Measurements of net CO_2_ assimilation (*A*_net_) and stomatal conductance (*g_s_*) were significantly higher in OQ414 compared to LH82 (independent *T-test* for each trait, *p* <0.05). All other steady state traits showed no significant difference between the two genotypes (*T-test, p* >0.05 (Fig. 6). *A/Ci* curves showed significantly higher rates of maximum carboxylation capacity of phosphoenolpyruvate (*V*_pmax_) in OQ414 compared to LH82, average *V*_pmax_ was 79.02±5.62 and 60.97±3.14 respectively (*T-test*, *p* <0.05) (Fig. 7). CO_2_ saturated photosynthetic capacity (*A*_sat_) was also significantly higher in OQ414 compared to LH82, average *A*_sat_ was 45.89±1.24 and 38.36±1.12 respectively (*T-test*, *p* <0.001) (Fig. 7).

**Figure 6:**
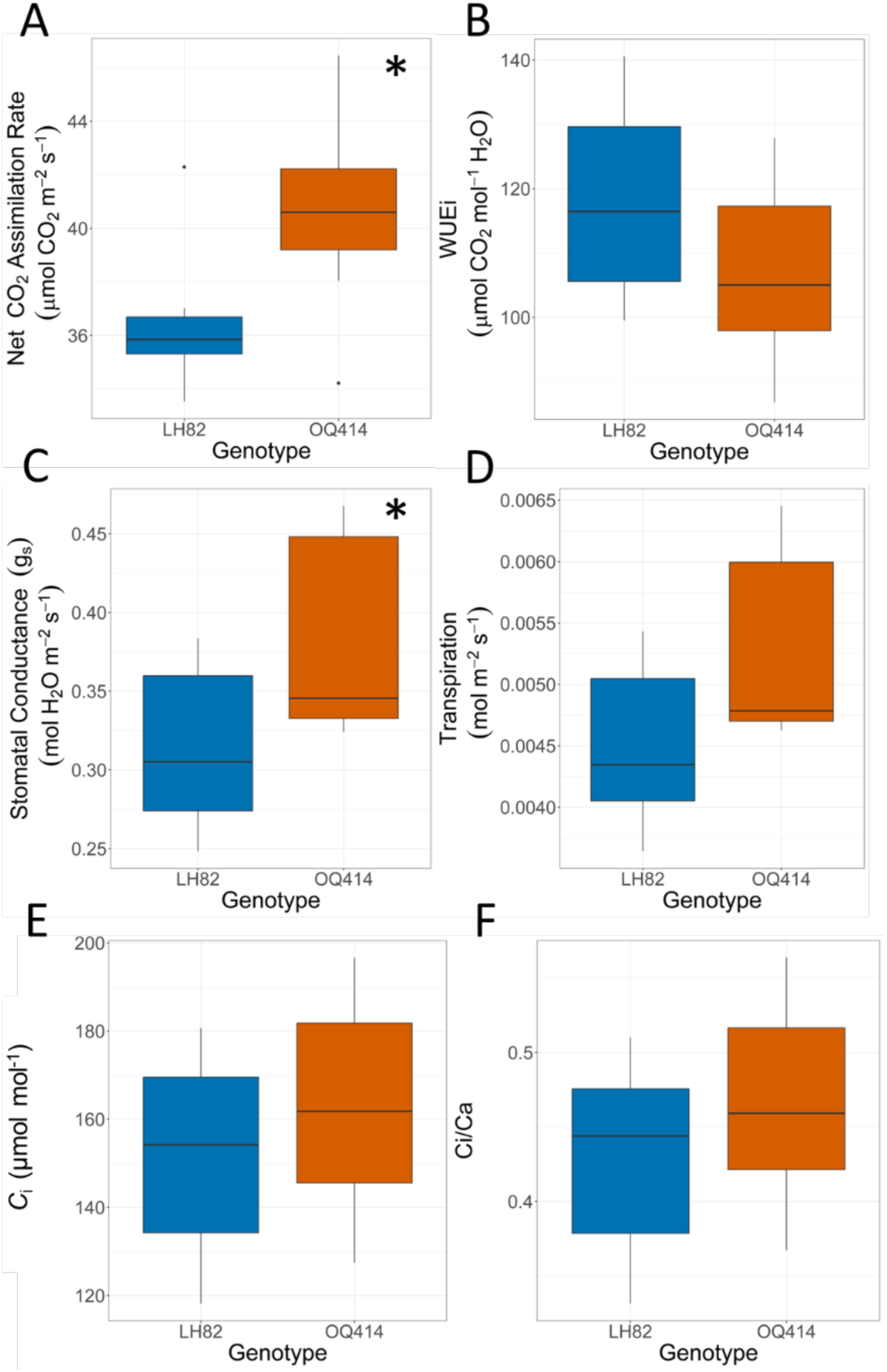
A) Net CO_2_ assimilation (*A*_net_), B) *WUE_i_* (*A/g_s_*), C) Stomatal conductance (*g_s_*), D) Transpiration (*E*), E) Intercellular CO_2_ (*C*_i_), and F) *C_i_/C_a_* under constant environmental conditions at a leaf temperature of 30°C, irradiance of 2,000 μmol PAR m^−2^ s^−1^, humidity of 70%, and atmospheric CO_2_ at 400 µmol mol^-^¹. Only *g_s_* ad *A_net_* were significantly higher in OQ414 compared to LH82 (*T-test*, *\** = *p*<0.05).

**Figure 7:**
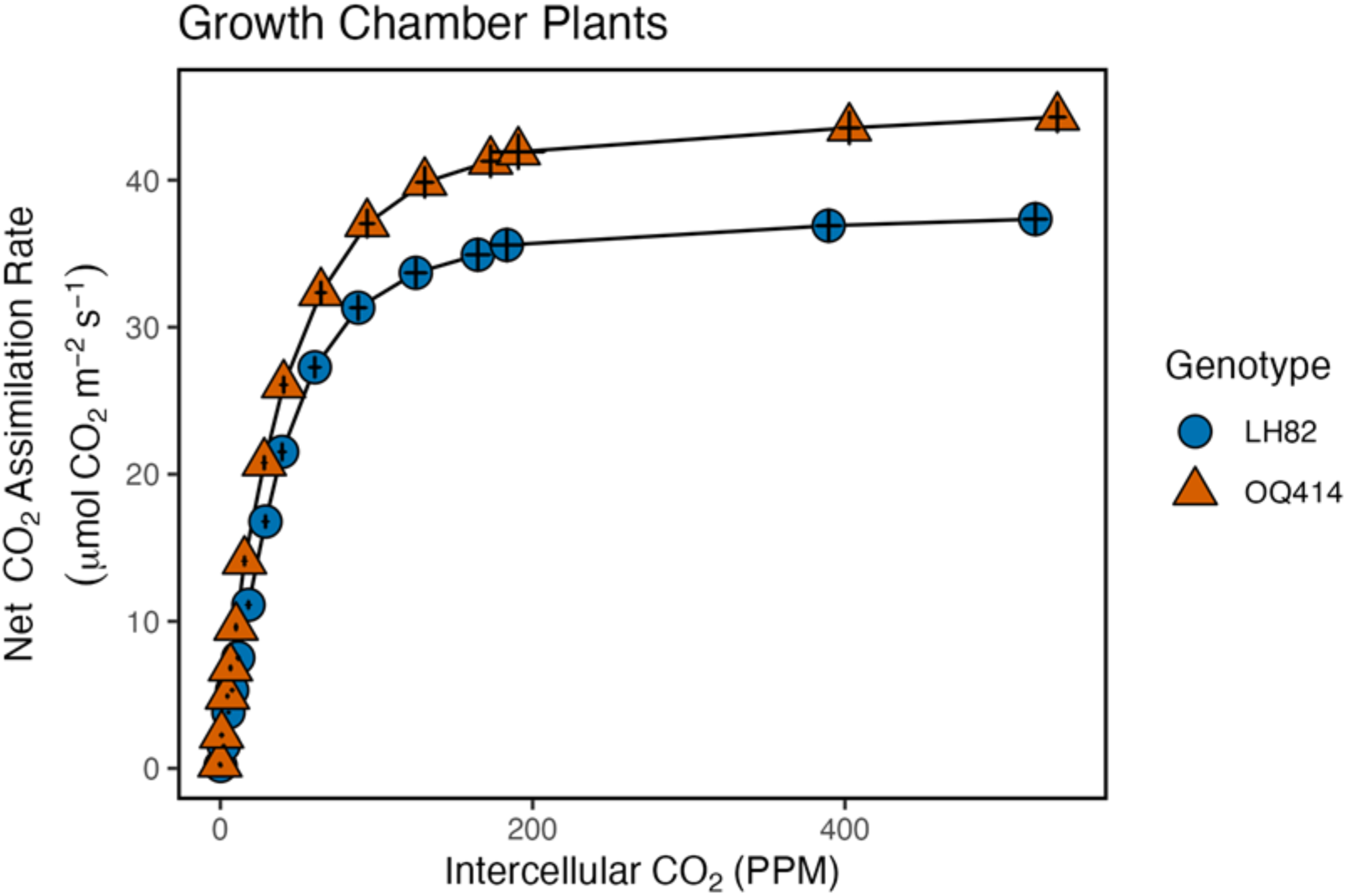
Net CO_2_ assimilation (*A*_net_) in response to changes in intercellular CO_2_ measured at a leaf temperature of 30°C, and irradiance of 2,000 μmol PAR m^−2^ s^−1^.

*In vitro* activity assays for CA, PEPC, and Rubisco were measured on the same leaves used for gas exchange. The first-order rate constant of CA (*k*_CA_; µmol CO_2_ m^−2^ s^−1^ Pa^−1^), PEPC (µmol HCO_3-_ m^−2^ s^−1^) activity, and Rubisco (µmol CO_2_ m^−2^ s^−1^) activity were all significantly greater in OQ414 compared to LH82 (Fig. 8).

**Figure 8:**
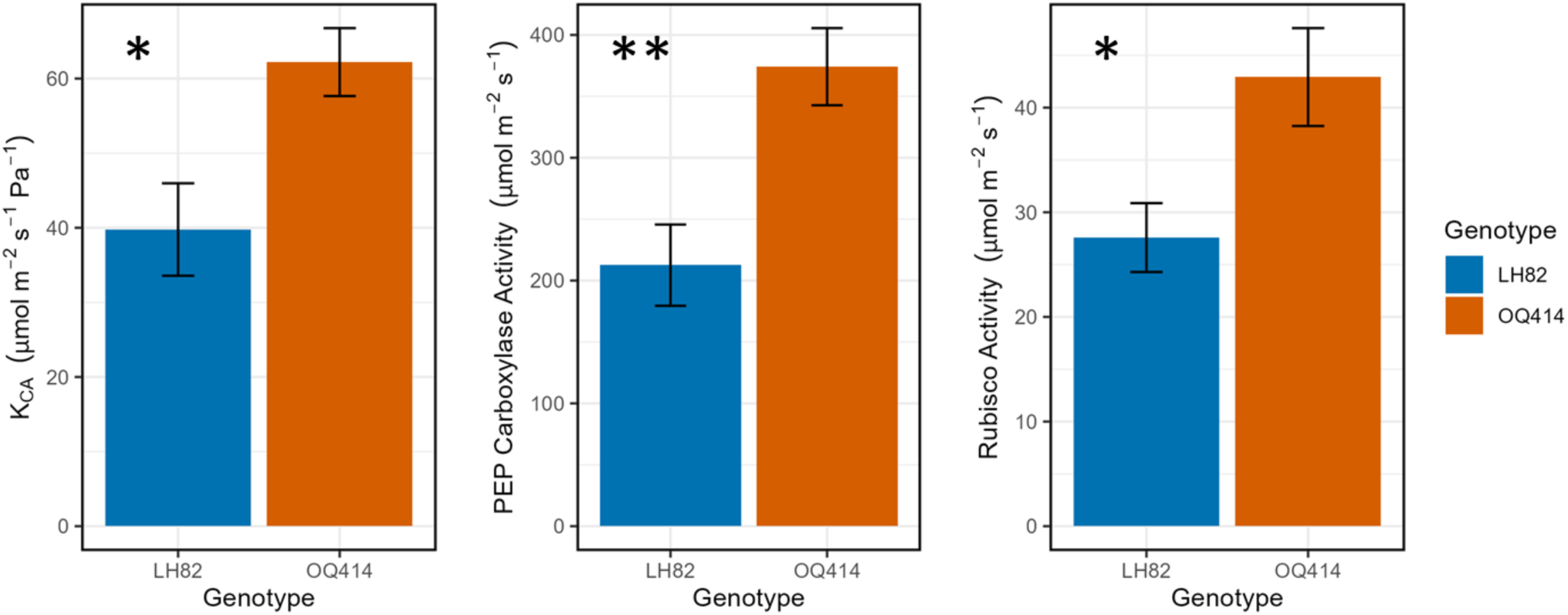
Carbonic Anhydrase (*k*_CA_), PEP Carboxylase, and Rubisco activity calculated for growth chamber grown plants. *k*_CA_ (*T-test*, *p* < 0.05, n = 5)*, PEPC (*T-test*, *p* < 0.01, n = 5)**, and Rubisco (*T-test*, *p* < 0.05, n = 5)* activity were all significantly higher in OQ414 (orange) compared to LH82 (blue). Bar plots represent average activity values and black error bars represent standard error.

## Discussion

Elucidating genetic regulators of δ^13^C_leaf_ in C_4_ species could allow for the broader implementation of this high-throughput proxy measure of *WUE_i_*. Here we identified a *Z. mays* line (OQ414) that contained a consistently negative δ^13^C_leaf_ value replicated in four field seasons. The average δ^13^C_leaf_ of OQ414 (-14.54±0.11‰) is significantly lower than other diverse *Z. mays* lines. Previously, a diverse set of 31 *Z. mays* inbred lines showed a δ^13^C_leaf_ range of -13.29 to - 11.61‰ (Twohey III et al., 2019). Using a QTL mapping approach, we identified a 5.1Mb region explaining 83.3% of the variation observed in δ^13^C_leaf_. This QTL region contains multiple *ca* genes already known to be implicated in CO_2_ fractionation. A complementation test and sequencing confirmed that a mutation in *cah1* underlies the observed δ^13^C_leaf_ value and led to the identification of the causative mutation.

Earlier studies across multiple species have observed more negative isotopic signatures when carbonic anhydrase is mutated. A *cah1 Dissociation* (*Ds*) transposon mutant in *Z. mays* reduced CA activity to 10% of the wildtype individual (Kolbe et al., 2018). Although the reduction in CA activity did not have a significant effect on photosynthetic rate at ambient CO_2_ concentrations, a decrease in δ^13^C_leaf_ was observed. Similar results have also been reported in *Flaveria bidentis*.

Antisense constructs targeted to mesophyll *cah1* reduced CA activity, resulting in reduced δ^13^C_leaf_ even when there was no reduction in net CO_2_ assimilation (Cousins et al., 2006). Thus, the observation in *Z. mays* that a decrease in CA activity also influences δ^13^C while maintaining photosynthetic rates at ambient CO_2_ is consistent with previous studies of *cah1* mutants in *F. bidentis*. However, to our knowledge this is the first case where a mutation in a carbonic anhydrase gene significantly reduces δ^13^C without a significant reduction in CA activity.

Based on the theory of carbon isotope discrimination, a significant reduction in δ^13^C could be explained in only a few ways. A reduction in δ^13^C can result from an increase in Φ (a decrease in the efficiency of CO_2_ concentration), an increase in the ratio of PEPC to CA activity (*V_p_/V_h_*), a decreased ratio of intercellular to ambient carbon dioxide concentrations (*C_i_/C_a_*), or a change in isotopic fractionation (Farquhar, 1983; Cousins et al., 2006, Studer et al., 2014). The significant increase in OQ414 PEPC activity and the corresponding greater net rates of CO_2_ assimilation would suggest that OQ414 has an equal or lower Φ compared to LH82. Therefore, a lower Φ in OQ414 would be expected to increase δ^13^C, opposite of what was observed. Although *k_CA_* and PEPC activities were significantly higher in OQ414, the *V_p_/V_h_* ratio for OQ414 (6.81±0.38) and LH82 (5.99±0.83) were not significantly different (*T-test, p* = 0.4, n = 5). In addition, no significant difference was observed in *C_i_/C_a_* (*T-test*, *p* = 0.3, n = 7). Thus, it is unlikely that an increase in *V_p_/V_h_* or decrease in *C_i_/C_a_* determine the more negative δ^13^C value observed in OQ414. These results uncouple the theoretical relationship between δ^13^C and *WUE_i_*. Although δ^13^C_leaf_ was significantly lower in OQ414 compared to LH82 we did not see a significant difference in *WUE_i_,* indicating that a change in isotope fractionation is likely driving δ^13^C_leaf_ without altering *g_s_* or *A_ne_*_t_. Rather than further elucidating the relationship between δ^13^C_leaf_ and *WUE_i_* in C_4_ species, this new finding of altered *cah1* fractionation adds a new consideration when using δ^13^C_leaf_ for breeding increased *WUE_i_*.

We have identified splice variants that encode for novel CA protein isoforms in OQ414 (Supp.7). These isoforms have altered domain structure and could affect the interaction between *cah1* and CO_2_. Given that the most common functional unit of β carbonic anhydrase is a tetramer, changes to domain structure and/or isoform length could impact the three-dimensional structure and catalytic activity of this key photosynthetic enzyme (Rowlett, 2010). Interestingly, we observe higher CA activity and photosynthetic rates in OQ414 relative to LH82, even though OQ414 carries a mutant allele of *cah1* that alters ^13^C discrimination, resulting in a more negative leaf δ^13^C value. This could be due to numerous differences between the genotypes, or perhaps, the novel CA isoforms may confer an advantage that increases rates of CO_2_ assimilation. Further studies are needed to investigate the fractionation of ^12^CO_2_ and ^13^CO_2_ by these novel CA protein isoforms.

Low atmospheric CO_2_ levels during the Oligocene epoch are believed to be the driver of C_4_ adaptation (Christin et al., 2008; Schlüter and Weber, 2020). Altering subcellular location and regulation of C_3_ *ca* genes allowed for the evolution of a carbon concentrating mechanism in C_4_ species, reducing the rate of photorespiration, and improving photosynthetic efficiency under low atmospheric CO_2_ levels (Ludwig, 2016; DiMario et al., 2017). Localization of *ca* has evolved from being chloroplastic in C_3_ species to primarily cytosolic in C_4_ species (Clayton et al., 2017). In addition to the initial relocation of CA in C_4_ plants, the gene structure of *ca* has continued to evolve within C_4_ species. Sequencing of tandemly duplicated *ca* genes of several monocot species showed fusion events and repeated duplication, as recently as after the divergence of sorghum and maize (Studer at al., 2016). Here we have isolated a small snapshot of *ca* evolution in *Z. mays* that appears to alter stable carbon isotope fractionation in a new way.

The inbred line OQ414 (Plant Variety Protection Application 009400244) was derived by Dow Elanco from the commercial maize hybrid PHI3540. Genotyping-by-sequencing data of OQ414 relative to a diversity panel of expired Plant Variety Protection (ExPVP) maize inbreds suggests that the parents of PHI3540 are PHG39 (PVP Application 008300115) and PHG50 (PVP Application 008300143), both developed by Pioneer Hi-Bred International (Studer, unpublished data). Neither PHG39 nor PHG50 carry the deletion allele identified in OQ414, indicating that this mutation arose spontaneously during the breeding cycle. Our result provide evidence that new isoforms encoding important C_4_ enzymes with altered biochemical properties can arise from spontaneous mutations, which ultimately can result in altered plant performance. Furthermore, this study highlights the lack of fundamental understanding about how δ^13^C is regulated in C_4_ plants and how it can be probed to advance understanding of crop performance.

## Supplementary Data

Table. S1. Primer Id and nucleotide sequences used to amplify and sequence DNA and cDNA fragments in OQ414 and LH82.

Table. S2. Data set containing markers and phenotypes used to QTL map δ^13^C_leaf_ and stomatal traits.

Fig. S3. Initial mapping to identify two significant δ^13^C_leaf_ QTL peaks on Chr. 3. Table. S4. Significant δ^13^C_leaf_ and stomatal QTL and their genomic positions.

File. S5. B73, OQ414, and LH82 *cah1* gDNA sequence. File. S6. B73, OQ414, and LH82 *cah1* cDNA sequence.

File. S7. B73, OQ414, and LH82 *cah1* predicted amino acid sequence.

## Acknowledgements

The authors would like to thank Lucas Roberts for identifying the OQ414 phenotype during field trials.

## Author Contributions

RJT: performed the F_2_ mapping population development, QTL mapping, complementation test, sequencing, and wrote the manuscript; JDC: performed the physiological gas exchange characterizations; MJR: sequenced the OQ414 deletion; JX : developed the model to extract and validate stomatal traits; ADBL: acquired funding, and supervised the research; ABC: acquired funding, and supervised the research AJS: conceptualized the project, acquired funding, and wrote the manuscript. All authors provided edits to the final manuscript.

## Conflict of Interest

The authors have no conflicts to declare.

## Funding

This work was supported by the United States Department of Agriculture—Hatch, a United States Department of Agriculture—Agriculture and Food Research Initiative grants (2019-67013- 29195) to AJS and ABC and (AFRI; 2020-67021-32799/project accession no.1024178) to ADBL from the USDA National Institute of Food and Agriculture, the National Science Foundation (grant no. PGR-1238030), and a Foundation for Food and Agriculture Research Graduate Student Fellowship.

## Data Availability

The corresponding authors will supply the data used in this manuscript upon request.

## Notes

### Competing Interest Statement

The authors have declared no competing interest.

